# Fast and flexible minimizer digestion with digest

**DOI:** 10.1101/2025.01.02.631161

**Authors:** Alan Zheng, Ishmeal Lee, Vikram S. Shivakumar, Omar Y. Ahmed, Ben Langmead

## Abstract

Minimizer digestion is an increasingly common component of bioinformatics tools, including tools for De Bruijn-Graph assembly and sequence classification. We describe a new open source tool and library to facilitate efficient digestion of genomic sequences. It can produce digests based on the related ideas of minimizers, modimizers or syncmers. `Digest` uses efficient data structures, scales well to many threads, and produces digests with expected spacings between digested elements. `Digest` is implemented in C++17 with a Python API, and is available open-source at https://github.com/VeryAmazed/digest.

## 1. Introduction

Digestion is the process of transforming a biological sequence into a shorter sequence that is still a useful reference for read alignment, sequence classification [Ahmed et al., 2023], or de novo assembly [Ekim et al., 2021]. Digestion works by selecting certain substrings to be kept according to strategies like minimizers [Roberts et al., 2004] or syncmers [Edgar, 2021]. The selected substrings are concatenated to form the digested sequence, which is often much shorter than the original. Digestion can be supplemented by “alphabet promotion” to further shorten the sequence and speed up matching algorithms [Ahmed et al., 2023, Ekim et al., 2021].

We present a new C++ software library called `digest` that performs digestion with (a) improved efficiency compared to previously described data-structures (b) efficient scaling to many threads, (c) three different substring-selection strategies: minimizers, syncmers or modimizers, and (d) an API allowing for various downstream uses, including Python bindings. In its command-line tool form, it can efficiently convert FASTA genomic sequences into digested sequences also in FASTA format.

We show that a combination of different data structures allow `digest` to work efficiently across a range of sequence lengths and window sizes. We show that naive approaches, as well as approaches based on the segment tree, are superior to approaches proposed in the past for common ranges of window sizes. Finally, we describe the API exposed by the `digest` tool and library, how it operates in parallel in its multithreaded mode.

## 2. Methods

`Digest` is a C++ software library that exposes an Application Programming Interface (API) for DNA sequence digestion. The following subsections detail its key data structures, how it was optimized, its interface, and how it maps to typical use cases.

`Digest` builds on the ntHash library Mohamadi et al. [2016], Kazemi et al. [2022] for efficient hashing of DNA sequences. Besides the features described in the following subsections, we describe additional implementation details in Supplementary Note S1.

### 2.1 Digestion schemes

`Digest` supports three strategies. The first uses “modimizers.” In this scheme, a length-*k* substring is included in the digest if and only if its hash value is equivalent to 0 mod *n*, where *k* and *n* are parameters. The second is based on “minimizers.” Here, a length-*k* substring is included in the digest if and only if its hash value is minimal in any of the length-(*w* + *k* − 1) substrings containing the *k*-mer. The third uses “syncmers.” In this scheme, a length-*w* substring is included in the digest if and only if the leftmost or rightmost *k*-mer (where *k < w*) of the substring has the minimal hash value among all the *w* − *k* + 1 length-*k* substrings.

In the event of multiple hash values that are both equal and minimal, we choose the rightmost by default.

### 2.2 Data structures

Modimizers are computationally easy to compute. But the other two schemes require the help of a data strucure to track hash values and compute minima. We implemented and benchmarked various structures supporting an *insert* query given an index and hash, as well as a *min* query which returns the index with the minimum hash in the current window. An index cannot be assumed to increase by one in each insert, as skips are possible due to unknown or ambiguous bases.

The <u>Naive</u> method uses a *deque* (double-ended queue), stored as an array in memory. *insert* simply adds an element at the head of the queue, simultaneously evicting an element from the tail. *Min* simply performs a linear scan of the queue to find the minimum element. The worst-case time is *O*(*nw*).

The <u>Naive-memo</u> method additionally *memoizes* (stores) the index of the minimum hash from the previous iteration. On an *insert* query, the new hash, if smaller than the stored minimum, replaces the memoized variable. If the stored minimum leaves the window, a linear scan is used to search for the new minimum. The *min* query, retrieves the memoized variable via a constant-time lookup (Supplementary Algorithm S2). Its worst-case time is *O*(*nw*), but its amortized cost is constant time, *O*(1) as we argue in Supplementary Note S2.

The <u>Set</u> method uses an ordered set, typically implemented as a red-black tree. We supplement the map with a deque of pointers to elements in the set. Inserting, removing, and finding the minimum element all take time that is logarithmic in the size of the window, with overall worst-case time therefore being *O*(*n* log *w*).

The <u>monotone</u> method uses a monotonic queue, which has been recommended for finding minimizers due to its linear runtime [Zheng et al., 2023]. It does indeed run in *O*(*n*). The queue holds the invariant that it is kept in increasing order. Old hashes are popped from the front as they leave the window. On an insertion, to maintain the invariant, all hashes that are greater than the inserted hash are deleted from the back. The hashes deleted from the back are of no use, since a smaller hash has entered the window. The minimum can be found by querying the front of the queue in *O*(1), while insertions can take *O*(*w*) time but amortizes to *O*(1).

The <u>segment tree</u> method uses a binary tree and maintains the invariant that a node holds the minimum of its two children. All leaves are situated at the same level, and the number of leaves is a power of two. If the specified window size does not give a power-of-two number of leaves, new leaves annotated with the maximum possible value are added as padding. The tree is represented implicitly in an array. The minimum can be queried in *O*(1) time by simply querying the root of the tree. Updates can traverse up the entire tree, taking *O*(log *w*) time (Supplementary Algorithm S3).

### 2.3. Multithreaded operation

We parallelize the digestion process by breaking the input sequence into partitions. We must allow for some overlap between the partitions for schemes that consider both large and small windows. A further complication is that the amount of overlap interacts with the treatment of non-ACGT characters, since the presence of ambiguous characters can effectively cause the large window to grow, so as to include the target number of non-ambiguous *k*-mers. This is discussed further in Supplementary Note S3.

### 2.4. Application Programming Interface

The `digest` software supports two APIs, one for C++17 and one for Python. The C++17 API is designed to be easy to use, with no dependencies on other libraries besides ntHash. Input sequence data is represented with an STL `string`, and results are appended to an STL `vector`. Users first instantiate the proper digester object from a hierarchy that includes an abstract parent class called `Digester`, a templated class called `WindowMin` for the windowed schemes, and a concrete class called `ModMin` implementing modimizers. For ease of use, we also provide a non-templated, concrete class called `Adaptive` that will attempt to select the optimal concrete class (and, therefore, data structure) for a given window-size scenario. `Adaptive64` is implemented alike with support for 64 bit hashes.

`digest` also has a Python API that applies the desired digestion scheme to a Python string, returning the result in a Python list. Note that this API wraps the `Adaptive` class, and so it allows the C++ library to choose the appropriate data structures according to the window-size scenario.

Example code using both APIs is included in the Supplementary Note S4.

## 3. Results

### 3.1 Data structures

The computational bottleneck to computing minimizers is the data structure used to facilitate both (a) the selection of the minimal value in a window (*min*), and (b) the updating of the window (*insert*). To understand the relative merits of the data structures, we implemented and conducted a benchmarking study wherein we applied all the data structures to an array of 10 million uniformly distributed hash values, executing the *insert* operation for every hash value and *min* operation for every window. As seen in Figure 1**A**, the best performing algorithms are <u>naive</u>, segment tree, and <u>naive-memo</u>. The <u>set</u> method’s runtime, which grows logarithmically with window size, exceeded the bounds of the chart and was omitted. <u>Monotone</u>, although a theoretically linear runtime data structure, also performed poorly, hampered by many conditional checks executed during the *insert* loop. Adaptive traces the shape of the best algorithms by determining the optimal data structure based on the desired window size. For small window sizes (6-14), <u>naive</u> outperforms all others due to a quick loop that is unrolled by the compiler and optimized with conditional move assembly instructions. For larger window sizes (*>*=16), <u>naive-memo</u> performs best by avoiding unnecessary window scanning while maintaining the simple loop structure of <u>naive</u>. In the final implementation of `digest`, we omitted the <u>monotone</u> and <u>set</u> methods, and suggest adaptive as the default back-end for general use cases.

**Fig. 1:**
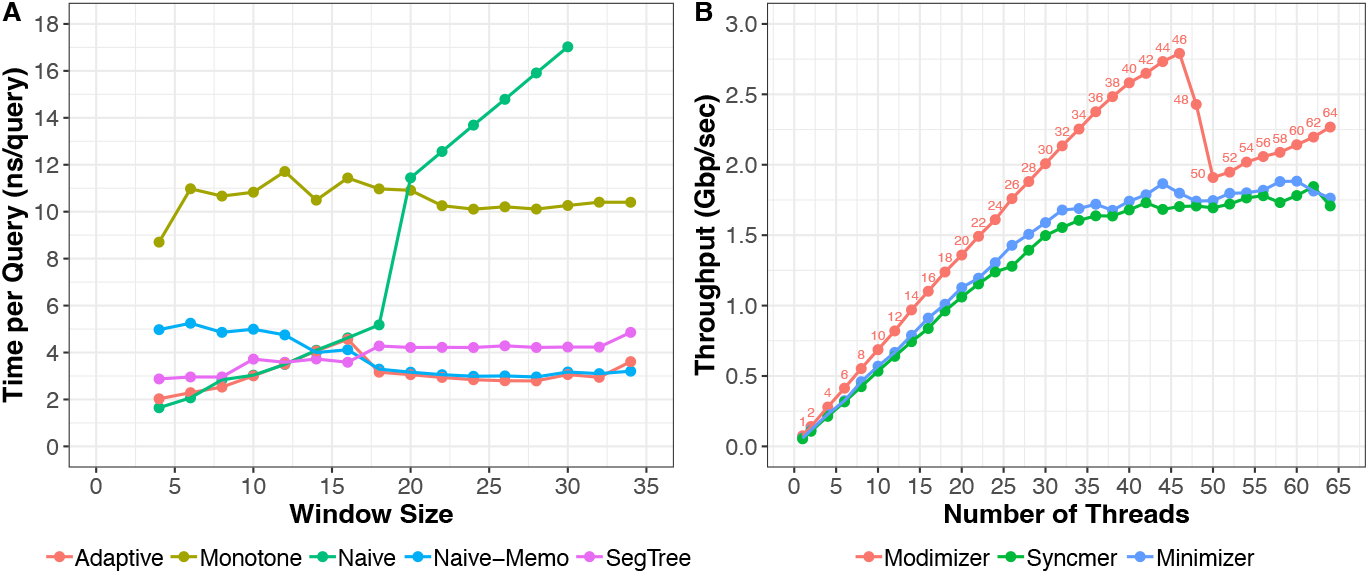
**(A)** Comparison of *min* query speed for different data structures as a function of window size. In this benchmark, each data-structure performs 10 million queries on an array of uniformly distributed 32-bit hash values. **(B)** Shows the throughput of the different digestion schemes in `Digest` (using segment tree data-structure) when computing the digest of a 62M human chromosome Y sequence consisting of only A/C/G/T characters. Benchmarking for both **(A)** and **(B)** were performed on a 48-core 3 GHz Intel Xeon Gold Cascade Lake 6248R CPU with 192GB RAM.

### 3.2. Thread scaling

To test the multi-threading scalability, we benchmarked the digest library when digesting the human chromosome Y sequence from T2T-HG002 assembly [Nurk et al., 2022] with an increasing number of threads.

Figure 1**B** shows the scalability of each digestion strategy. All three strategies scale linearly with an increasing number of threads until we approach the number of cores on our machine, which was 48. Around 48 threads, we observed a decline in throughput for modimizers, and a less pronounced slowdown for syncmers and minimizers. The throughput observed as we continue to increase the number of threads (*>* 48) can be attributed to better load balancing for this machine.

As expected, the modimizer scheme has the highest throughput given its simpler strategy that do not require an auxilary for range-minimum queries. Overall, the `digest` library shows consistent throughput improvements for each digestion scheme as we add more and more threads reaching speeds ranging from 1.7 Gbps to 2.8 Gbps.

## 4. Discussion

We present a new, efficient C++/Python software library called `digest` implementing modimizers, minimizers and syncmers. We benchmarked the tool comprehensively for multi-thread scalability and across different back-end implementations.

We identified various avenues for future development. `digest`’s thread scaling capability is currently limited when dealing with non-ACGT characters, and additional policies for handling a general input alphabet is warranted for broader applicability beyond biological sequence data.

Additionally, our multi-threading strategy divides the input into a number of equal-sized, overlapping partitions, where the number of partitions equals the number of simultaneous threads. In the future, we plan to implement a strategy that uses a work queue so that the number of threads can be specified independently of the size of each individual partition, to allow for better load balance at a smaller numbers of threads.

Since one of the major uses of minimizer digestion is in settings where the digested sequence should undergo “alphabet promotion,” a goal for future versions of the `digest` library will be to support this as an immediate output of the digestion process.

For instance, users could digest a long biological sequence into a shorter “promoted” sequence of, say, 8-bit minimizer symbols, in a single API call.

Lastly, the current field of minimizer research is constantly evolving with a great interest in deriving schemes with lower and lower expected densities [Koerkamp and Pibiri, 2024, Kille et al., 2024]. The closer the density is to the theoretical lower bound, the smaller the “digest” will be which typically translates into both reduced running time and memory costs downstream. It will be important in the future for new schemes to be added into the digest library in order to provide users with different options since different schemes can perform optimally in different use-cases.

## 5. Author contributions and Competing interests

A.Z. and I.L. implemented the software package. V.S.S and O.Y.A contributed to the software development. A.Z., I.L., and O.Y.A benchmarked the software. All authors contributed to and reviewed the manuscript. No competing interest is declared.

## 6. Acknowledgments

We thank Ragnar Groot Koerkamp for his comments on the manuscript.

## 7. Funding

V.S.S, O.Y.A, and B.L. were supported by NIH grant R35GM139602 to B.L.

## Notes

### Competing Interest Statement

The authors have declared no competing interest.

