## Supplement for "Fast and flexible minimizer digestion with digest"

#### Algorithms

In this section, we include the pseudocode for the different data-structures used within the `digest` library. For each data-structure, we include the function to insert a new integer value as well as the function to retrieve the smallest integer value (i.e. minimizer).

##### S1 Naive

**Algorithm 1** Naive Insert Function

**Require:** $hash$: hash value

$arr[i] \leftarrow hash$

$i \leftarrow (i + 1) \bmod w$ ▷ Updates next position for update, it is a circular array

**Algorithm 2** Naive Min Function

**Require:** $arr$: array of last $w$ hash values

$i \leftarrow 0$

**for** $j = 1$ to $w - 1$ **do** ▷ Loop through and find minimum

  **if** $arr[j] < arr[i]$ **then**

    $i \leftarrow j$

  **end if**

**end for**

**return** $arr[i]$

#### S2 Naive-Memo

**Algorithm 3** Naive-Memo Insert Function

**Require:** $hash$: hash value, $last$: index of current minimum hash value
$arr[i] \leftarrow hash$
**if** $arr[i] < arr[last]$ **then** ▷ Update memo-ized value
    $last \leftarrow i$
**else if** $last = i$ **then** ▷ When minimum leaves the window, rescan
    **for** $j = 0$ to $w - 1$ **do**
        **if** $arr[j] < arr[last]$ **then**
            $last \leftarrow j$
        **end if**
    **end for**
**end if**
$i \leftarrow (i + 1) \bmod w$ ▷ Updates next position for update, it is a circular array
**Ensure:** $last$: updated index of minimum hash value in $arr$

**Algorithm 4** Naive-Memo Min Function

**Require:** $arr$: array of last $w$ hash values, $last$: index of minimum hash value
**return** $arr[last]$

#### S3 Segment tree

**Algorithm 5** Segment Tree Insert Function

**Require:** $hash$: hash value, $w$: number of hash values in data structure
$segtree[i] \leftarrow hash$
**for** $level = 0$ to $\lceil \log_2(w) \rceil$ **do** ▷ Go through each level of segment tree
    $segtree[i/2] \leftarrow \max(segtree[i], segtree[i \oplus 1])$ ▷ XOR gets sibling in tree
    $i \leftarrow i/2$
**end for**

**Algorithm 6** Segment Tree Min Function

**Require:** $segtree$: segment tree of $w$ hash values

```
return segtree[1]
```

#### S4 Adaptive

This data-structure tries to combine the optimal performance of the Naive and Naive-Memo approaches in different k-mer ranges. Therefore, in each function below, you will observe an if statement that checks what $k$ is being used in order to choose the appropriate function.

**Algorithm 7** Adaptive Insert Function

**Require:** $hash$: hash value, $w$: number of hash values in data structure

```
if w < 16 then
    naive_insert_function(hash)
else
    naive_memo_insert_function(hash)
end if
```

**Algorithm 8** Adaptive Min Function

**Require:** $w$: number of hash values in data structure

```
if w < 16 then
    naive_min_function(hash)
else
    naive_memo_min_function(hash)
end if
```

### Notes

#### S1 Design details

- **Deques**: Whenever a "deque" is mentioned, it is actually implemented using a static circular array.
- **Templating**: With the use of templating in C++, the compiler knows the size of the large window size at compile time, allowing the compiler to choose optimizations such as loop unrolling.
  - This helps to explain the gap in performance between Adaptive and templated data structures.
- **Bit-packing of hash and index**: In `digest`, 32-bit index and 32-bit hash are packed together into 64-bit words which allows for many optimizations such as better memory locality.
  - Comparisons and moves can be performed in one computer instruction instead of two Thus, Adaptive-64 can perform much worse (2x).
- **Handling of non-ACGT characters**: There are currently two policies user can choose from when handling non-ACTG characters.
  - **SKIPOVER**: Every k-mer that contains a non-ACTG character is skipped
  - **WRITEOVER**: Replaces every non-ACTG character with 'A'

#### S2 Amortized Time Analysis of Naive-Memo Algorithm

- **Worse Case**: Its worst-case time is $O(nw)$, but for a random hash, an amortized complexity analysis can be performed.
- **Amortized Analysis:**
  - **Step 1**: Begin by reading in the first $w$ hash values and find the smallest value, therefore the time complexity is $O(w)$
  - **Step 2**: Each time we move the sliding window by 1, there are two different types of operations we can perform:
    - **Find a value smaller than current $w$ values**: Once we find a smaller value, we will ultimately have to perform $O(w)$ scan once that value falls out of the window to find the new minimum.
    - **Encounter a value larger than current minimum**: Therefore, we can quickly return the memo-ized minimum in constant time, $O(1)$

- **Step 3**: Compute the time complexity spent in each type of step:
  * **Find a value smaller than current $w$ values**: The probability of finding a value smaller than current window is $\frac{1}{w+1}$ and there are $n-w$ opportunities. Therefore, the total time spent in this step is $\frac{1}{w+1} * (n-w) * w$
  * **Encounter a value larger than current minimum**: The probability of finding a value larger than the current minimum is $\frac{w}{w+1}$ and there are $n-w$ opportunities, therefore, the total time cost is $\frac{w}{w+1} * (n-w)$
- **Step 4**: Compute total time complexity and amortized time:
  * Total time: $\frac{2w}{w+1} * (n-w) = O(n)$
  * Amortized time: $\frac{O(n)}{n} = O(1)$

#### S3 Multi-threaded Mode

- **Proper handling of digestion**: For schemes that use a large window, we want every large window to be considered exactly once, and for those that don't, we want each kmer to be considered exactly once.

- **Logic used to separate windows:** For kmers, if the the $i$-th kmer ends at index $j$, then the $i+1$-th kmer should should start at index $j-k+1$. So then if segments A and B are two adjacent segments in the original sequence and segment A ends at index $j$, then segment B should started at $j-k+1$.

  - Similar logic can be applied when ensuring each large window is only considered once. This method of threading, although very straight forward to implement is likely not the most efficient implementation.

- **Issue of non-ACTG characters**: You can only guarantee that each kmer/large window is considered exactly once if you do not skip over any kmers for containing non-ACTG characters, i.e. either there are no non-ACTG characters, or you are using the skip-over policy.

- **Example**: $seq$ = ACTGANACNACTGA, $k$ = 4, $w$ = 4, # of threads = 2

  - There are only 4 valid k-mers in this sequence
  - Therefore, only 1 valid large window
  - But we can't know this until it actually goes through the sequence, so it's going to try to partition the sequence into ACTGANACNA, and ANACNACTGA which now each have 0 valid large windows.

- **Using WRITEOVER can get past the issue**: As mentioned in the Design Details section, even if your sequence has non-ACTG characters in it, you can still perform multi-threading if you use the write-over policy as it replaces every non-ACTG character it encounters with an A.

#### S4 Application Programming Interfaces (APIs)

The `digest` library supports two different APIs in order to be used by users. The sections below detail how `digest` library can be utilized as Python library or as C++ library. Both examples focus on the *minimizer* scheme however similar approaches can be used for *modimizers* and *syncmers*.

##### S4.1 Python Bindings

```
>>> from Digest import window_minimizer
>>> window_minimizer("GATATAATAGACTATAACTATAGATACTA", k=15, w=5, include_hash=True)
[(1, 41488463), (2, 740555263), (4, 2103507089), (8, 724537772), (9, 2140030416), (14, 1405637373)]
>>> window_minimizer("GATATAATAGACTATAACTATAGATACTA", k=15, w=5, include_hash=False)
[(1, 41488463), (2, 740555263), (4, 2103507089), (8, 724537772), (9, 2140030416), (14, 1405637373)]
[1, 2, 4, 8, 9, 14]
```

##### S4.2 C++ library

```
#include "digest/digester.hpp"
#include "digest/window_minimizer.hpp"

int main() {
    std::string dna {"ACGATAATAGACTATAACTATAGATACTA"};
    std::vector<std::pair<uint32_t, uint32_t>> output;
    int k = 15; int w = 7;

    digest::WindowMin<digest::BadCharPolicy::WRITEOVER, digest::ds::Adaptive> digester(dna, k, w);

    digest.roll_minimizer(dna.length(), output);
    # output vector will contain pairs of (position, hash value)

}
```

##### S5 Density

Expected densities where $w$ is the number of k-mers in the large window:

- **Minimizer**: $\frac{2}{w+1}$
- **Syncmers**: $\frac{2}{w}$
- **Modimizers**: Can be represented as a Bernoulli($\frac{1}{M}$) so its expected density is $\frac{1}{M}$

In the figure section below, we include empirical data showing that `digest` library generates minimizer vectors closely following the expected densities.

#### Figures

- For the density experiments, multiple long strings of ACTG characters are randomly generated and used as input for different digestions schemes to compute the observed density.
- Experimental code can be found here: `https://github.com/VeryAmazed/digest/blob/main/tests/density/Results.md`

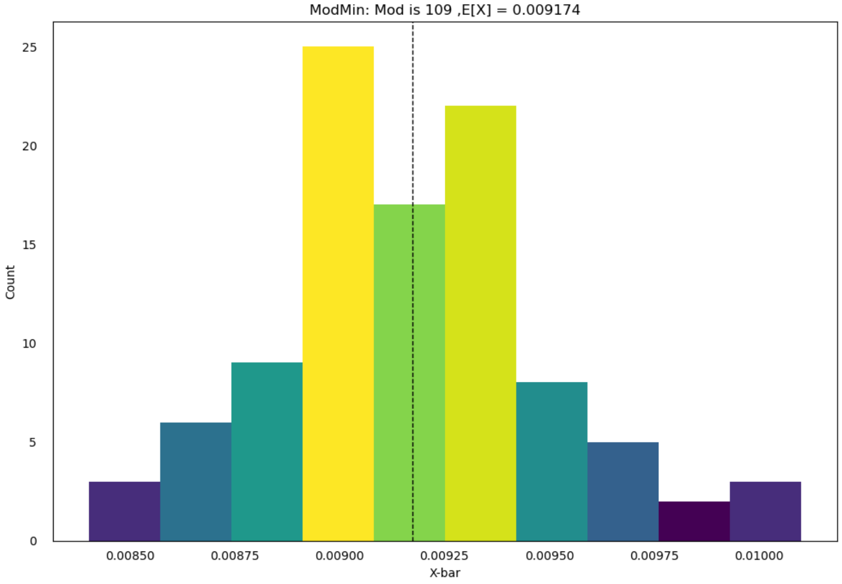


Figure S1: Average density of the modimizers obtained over many random trials with modulus 109. Dotted line shows expected density based on theory.

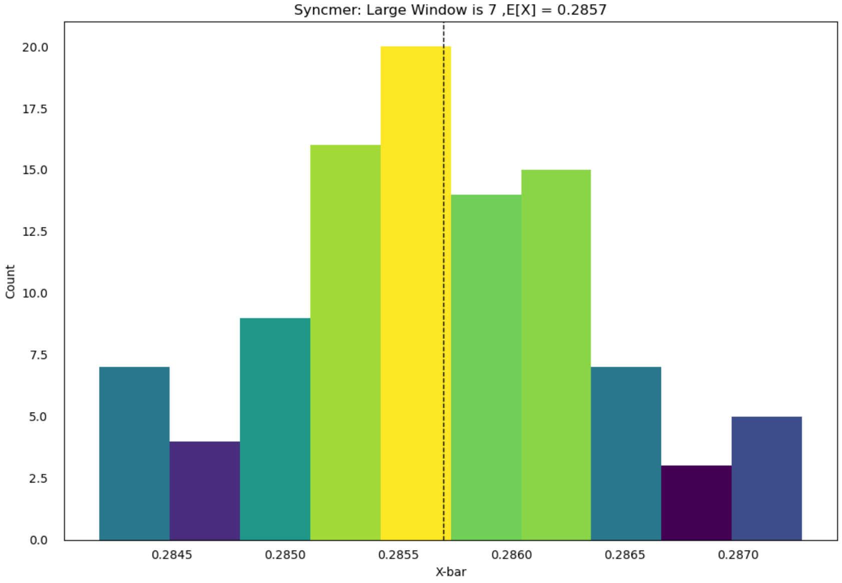


Figure S2: Average density of the syncmers (w=7) obtained over many random trials. Dotted line shows expected density based on theory.

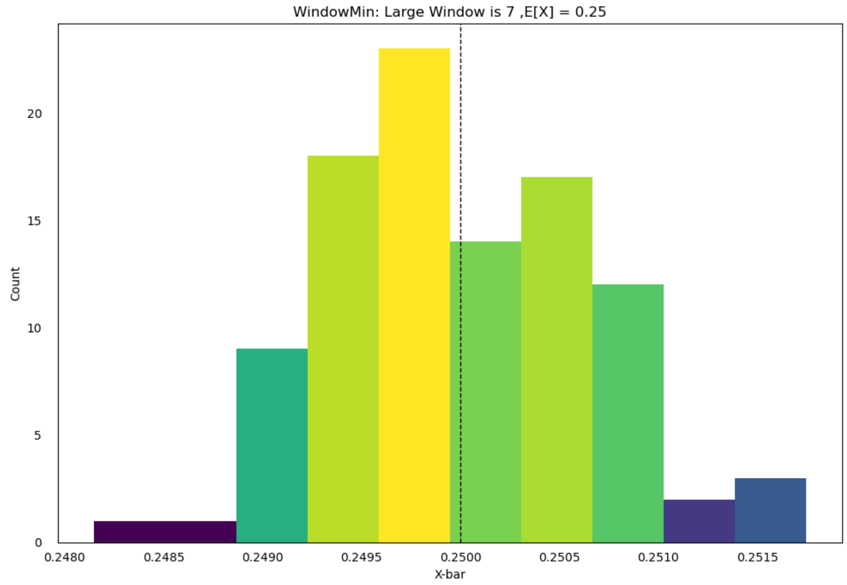


Figure S3: Average density of the minimizers (w=7) obtained over many random trials. Dotted line shows expected density based on theory.
